# A missense mutation of *Ip3r1* in *Dp2* mice leads to short-term mydriasis and unfolded protein response in the iris constrictor muscles

**DOI:** 10.1101/465591

**Authors:** Bing Chen, Chongyang Qi, Li Chen, Mengjun Dai, Yayou Miao, Rui Chen, Wane Wei, Shun Yang, Hongling Wang, Xiaoge Duan, Minwei Gong, Wang Yi, Zhengfeng Xue

## Abstract

*Ip3r1* encodes an inositol 1,4,5-triphosphate-responsive calcium channel. Mutations in the *Ip3r1* gene in humans may cause Gillespie syndrome (GS) typically presents as fixed dilated pupils in affected infants, which was referred to as iris hypoplasia. However, there is no report of mice with *Ip3r1* heterozygous mutations showing dilated pupils. Here, we report a new *Ip3r1* allele (dilated pupil 2; *Dp2*) with short-term dilated pupil phenotype derived from an N-ethyl-N-nitrosourea (ENU) mutagenesis screen. This allele carries a G5927A transition mutation, which is predicted to result in a C1976Y amino acid change in the open reading frame. Histology and pharmacological tests show that the dilated pupil phenotype is a mydriasis caused by the functional defect in the iris constrictor muscles in *Dp2*. The dilated pupil phenotype in *Dp2* was referred to as mydriasis and excluding iris hypoplasia. IHC analysis revealed increased expression of BIP protein, the master regulator of unfolded protein response (UPR) signaling, in *Dp2* mice that did not recover. Apart from the dilated pupil phenotype (mydriasis), there are no other abnormal phenotypes including *Ip3r1*-related ataxia that may be found. This study is the first report of an *Ip3r1* mutation being associated with the mydriasis phenotype. *Dp2* mice represent a valuable self-healing model that may be used to study the therapeutic approach for *Ip3r1*-related diseases or diseases caused by similar pathomechanisms.

## INTRODUCTION

Inositol 1,4,5-trisphosphate receptors (IP3Rs) are ligand-gated ion channels that release calcium ions (Ca^2+^) from the endosarcoplasmic reticulum (ER) to the cytoplasm in response to the binding of inositol 1,4,5-trisphosphate (IP3) (Berridge et al., 2003; Berridge et al., 2000). Three subtypes of this receptor have been identified, namely, IP3R1, IP3R2, and IP3R3 (also known as ITPR1, ITPR2, and ITPR3, respectively). These three isoforms share a common structure consisting of an IP3-binding core in the N-terminal portion, a channel domain at the extreme C-terminal end and a central coupling or modulatory domain (Foskett et al., 2007; Hamada et al., 2014; Yule et al., 2010). These subtypes are co-expressed in a variety of cells and can form either homotetramer or heterotetramer Ca^2+^ release channels with each having distinct channel properties (Hours and Mery, 2010).

*Ip3r1* is predominantly expressed in the central nervous system (CNS), especially in cerebellar Purkinje cells (PCs) (Furuichi et al., 1993). *Ip3r1* gene mutations in humans are associated with different types of autosomal dominant spinocerebellar ataxia (SCA) including late-onset spinocerebellar ataxia type 15 (SCA15) (Hara et al., 2008; Marelli et al., 2011; Obayashi et al., 2012; Tipton et al., 2017; van de Leemput et al., 2007; van Dijk et al., 2017), congenital nonprogressive spinocerebellar ataxia and mild cognitive impairment (SCA29) (Huang et al., 2012; Klar et al., 2017; Wang et al., 2018; Zambonin et al., 2017), infantile-onset cerebellar ataxia with mild cognitive deficit (Sasaki et al., 2015), and childhood-onset ataxic cerebellar palsy with moderate intellectual disability (Parolin Schnekenberg et al., 2015). Recently, mutations in the *Ip3r1* gene were identified in patients with Gillespie syndrome. Gillespie syndrome typically presents as fixed dilated pupils (referred to as iris hypoplasia) in affected infants. The key extra-ocular features of Gillespie syndrome are congenital hypotonia, non-progressive cerebellar hypoplasia, ataxia and variable, usually mild, neurocognitive impairment. However, no major abnormality in eye development has been described in mice with *Ip3r1* heterozygous null mutations (De Silva et al., 2018; Dentici et al., 2017; Gerber et al., 2016; McEntagart et al., 2016).

We have previously shown that an ENU-induced mutation in the *Nrg1* gene leads to the dilated pupil phenotype in *Dp1* (Dilated pupil mutation 1) mice (Chen et al., 2011). Another short-term dilated pupil phenotype mice *Dp2* (Dilated pupil mutation 2) is also reported in this study. Here, we report that the abnormal phenotype is due to a mutation in the *Ip3r1* gene, which causes a functional defect in the iris constrictor muscles. Apart from the dilated pupil phenotype (mydriasis), there are no other abnormal phenotypes including ataxia that may be found.

## RESULTS

### Generating a mutant mouse with short-term dilated pupil phenotype using ENU mutagenesis

In a dominant ENU mutagenesis screen for abnormal eyes, we identified a mouse mutation *Dp2* with the bilaterally dilated pupil phenotype at 2-week time point (Fig. 1A). The ENU-mutagenesis parental strain was C57BL/6 (B6) male mice, and this was crossed with untreated B6 female mice for screening. After mating the *Dp2* mice with wild-type B6 mice, a ratio of 359 of the 770 progeny were recorded to have the dilated pupil phenotype. Interestingly, all pupils of *Dp2* recovered and were basically back to normal size after 4 weeks (data not shown).

**Figure 1.**
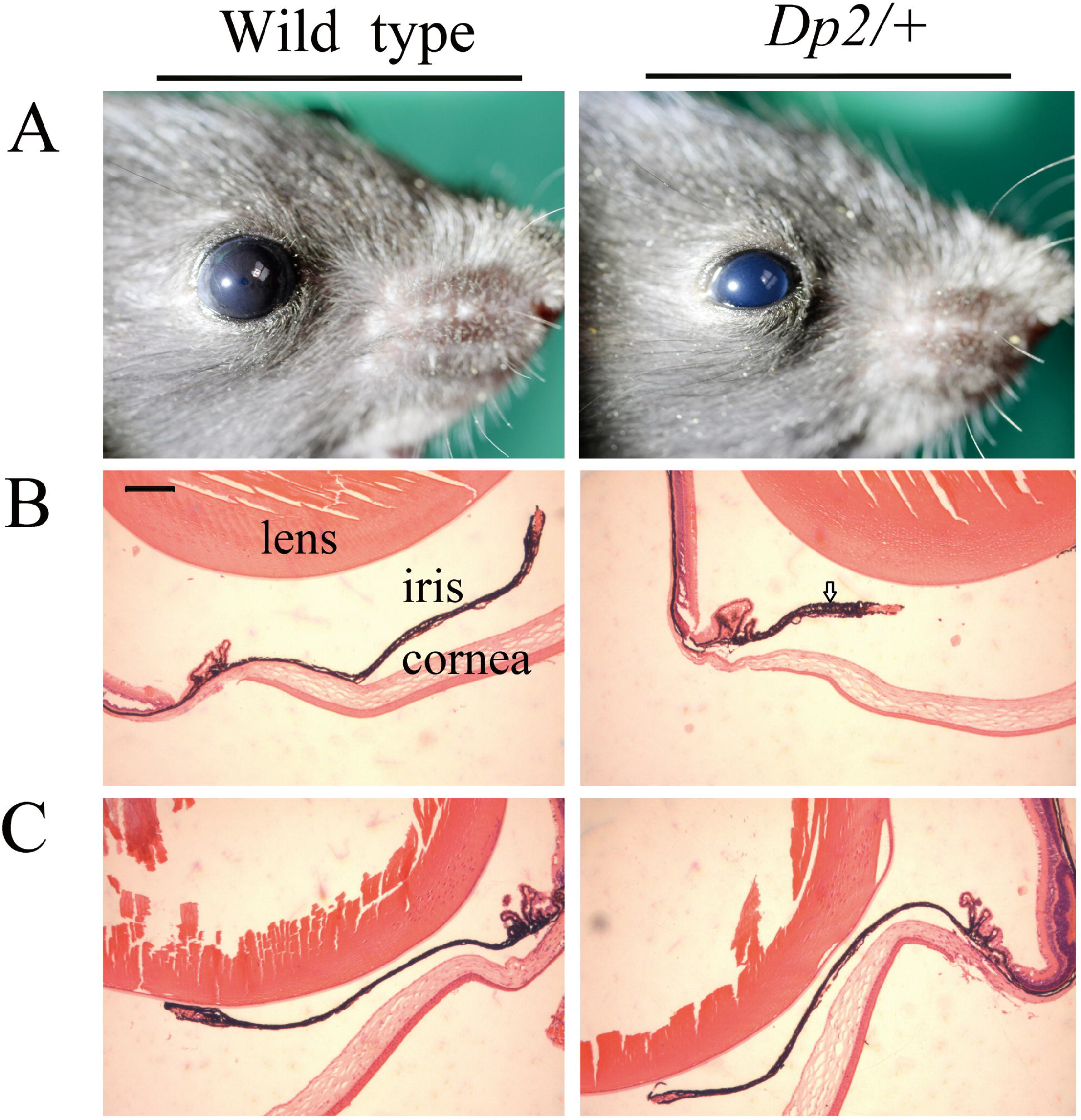
Short-term mydriasis phenotype in *Dp2* mice. (A)Wild-type mouse with normal pupil and *Dp2* heterozygous mouse with a dilated pupil phenotype. (B, C) Eye pictures of hematoxylin and eosin staining from wild-type and *Dp2* mice. *Dp2* mice that did not recover and littermate controls (B). *Dp2* mice that were recovered and littermate controls (C). The arrow in (B) shows the short and unstretched iris in *Dp2* mice that did not recover. At least three mice were used for each analysis, and the representative data were shown. Scale bar=100 µm.

### The *Dp2* phenotype is the result of a functional defect of the iris constrictor muscles

Hematoxylin and eosin staining showed *Dp2* mutants that did not recover to have intact but short and unstretched irises, compared with littermate control mice and *Dp2* mice that were recovered (Fig. 1B, C). The results indicate that the defect in *Dp2* mutants affects the pathway for the pupillary light response or in the muscles of the iris. The dilated pupil phenotype in *Dp2* was therefore referred to as mydriasis, thereby excluding iris hypoplasia. To further analyze the defect, a 1% pilocarpine solution (a nonselective muscarinic cholinergic receptor agonist) was applied as drops in the eyes of *Dp2* mice. Consequently, the mutant pupils fail to constrict indicating that the defect in *Dp2* mutants is in the iris constrictor muscles.

### The *Dp2* phenotype is caused by a point mutation in the *Ip3r1* gene

For initial mapping, we tested genomic DNA from 50 N2 samples with microsatellite markers across the whole genome. The mutation was mapped to chromosome 6 and had 1 exchange with marker *D6Mit230*, which was located at 45.74cM. In order to further refine the map position, genomic DNA from 400 N2 mutants was analyzed using microsatellite markers across the whole genome. The mutation was mapped to a 4.78Mb region on chromosome 6, between microsatellite *D6Mit149* and *D6Mit148* (Fig. 2A). The region contained 10 protein coding genes (*Arl8b*, *Bhlhe40*, *Crbn*, *Edem1*, *Il5ra*, *Ip3r1*, *Lrrn1*, *Setmar*, *Sumf1* and *Trnt1*), 2 unclassified non-coding RNA genes (0610040F04Rik and Gm17055). Sequence analysis of these genes revealed no apparent nucleotide changes in *Dp2* mice except fo*r Ip3r1*.

**Figure 2.**
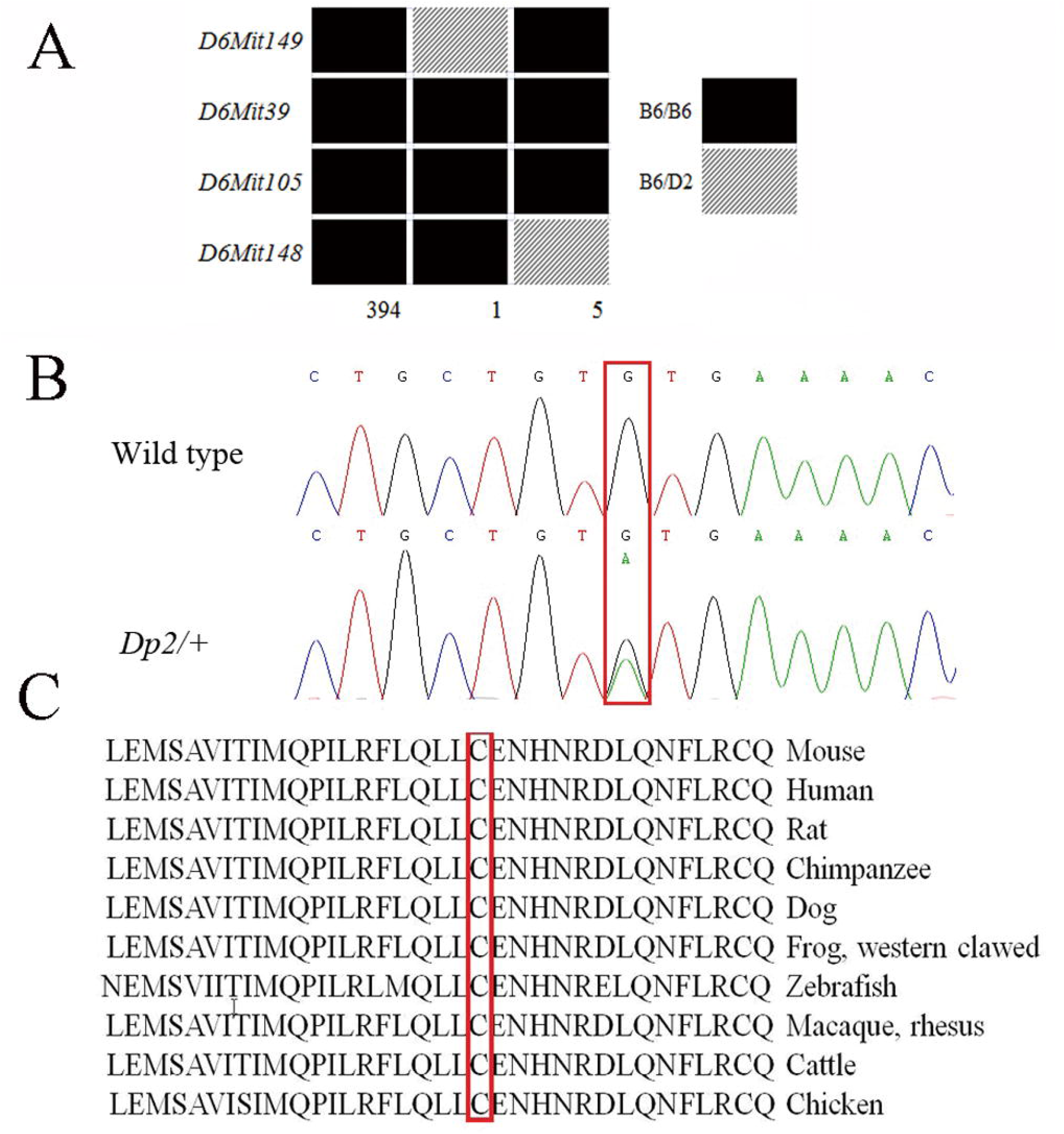
*Dp2* mapping and mutation analysis. (A) Genetic mapping places the mutant in the region between markers *D6Mit149* and *D6Mit148*. (B) Sequence analysis of the *Itpr1* gene showed a G to A transition mutation, which resulted in a C1976Y amino acid change in the open reading frame in *Dp2* heterozygous mice compared to wild-type B6 mice. (C) Sequence alignment across multiple species revealed that this cysteine acid residue is highly conserved among vertebrates.

In the *Ip3r1* gene, we identified a G5927A transition mutation in exon 46 (Fig. 2B), which is predicted to result in a C1976Y amino acid change in the open reading frame. Sequence analysis of three different wild-type strains (C3H/He, 129 and DBA/2) excluded a general polymorphism at this site (data not shown). Sequence alignment across multiple species revealed that Cys-1976 is a highly evolutionarily conserved amino acid in IP3R1 suggesting conservation of function (Fig. 2C).

### UPR activation in the iris constrictor muscles of unrecovered *Dp2* mice

To test whether the *Ip3r1 ^C1976Y^* mutation leads to alterations of IP3R1 protein levels in the iris constrictor muscles of *Dp2* mice, we performed immunohistochemistry (IHC) tests on eye sections from *Dp2* mice that were recovered and those that did not recover. The results revealed that IP3R1 protein expression was detected and similar in iris constrictor muscles of all mice (Fig. 3A, B). Mutant proteins within both the cytoplasm and the ER stimulate the unfolded protein response (UPR). An ER chaperone, BIP acts as the master regulator of UPR signaling (Hendershot, 2004). IHC analysis using anti-BIP antibody showed that, compared with wild-type littermates, BIP levels were increased in *Dp2* mice that did not recover (Fig. 3C). There was no significant difference in recovered *Dp2* mice and wild-type littermates (Fig. 3D). Failure of the UPR to resolve ER stress results in the stimulation of caspase-mediated apoptotic pathways (Allen et al., 2016; Kim et al., 2008). IHC analysis however, showed no significant apoptotic caspase 3 cells observed in the iris constrictor muscle of all mice (data not shown).

**Figure 3.**
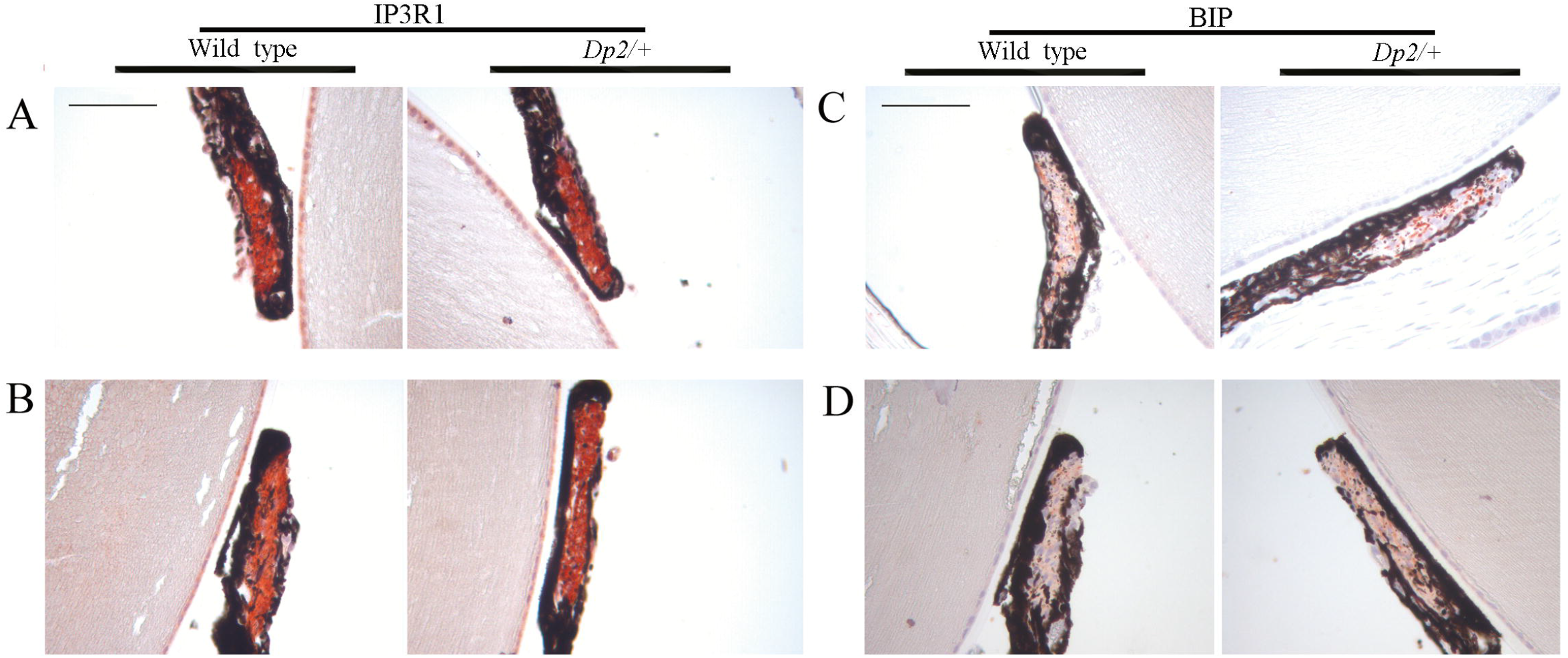
IHC analysis of iris constrictor muscle using anti-IP3R1 and anti-BIP antibodies. (A, B) IHC analysis of iris constrictor muscle using anti-IP3R1 antibody. *Dp2* mice that did not recover and littermate controls (A). *Dp2* mice that were recovered and littermate controls (B). The results showed similar IP3R1 protein level in the iris constrictor muscles between *Dp2* mice and wild-type littermates. (C, D) IHC analysis of iris constrictor muscle using anti-BIP antibody. *Dp2* mice that did not recover and littermate controls (C). *Dp2* mice that were recovered and littermate controls (D). IHC results revealed that the expression of BIP was upregulated in iris constrictor muscles of *Dp2* mice that did not recover. The expressions of IP3R1 and BIP are shown in red. At least three mice were used for each analysis, and the representative data were shown. Scale bar=100 µm.

### *Dp2* mutants showed no obvious cerebellar ataxia symptom

Apart from the dilated pupil phenotype (mydriasis), there are no other abnormal phenotypes including ataxia that may be found. Cerebellar size of *Dp2* was comparable to wild-type littermates (data not shown). The expression level of IP3R1 in the cerebellum of *Dp2* mice was equivalent to that of wild-type littermates (Fig. 4A, B). IHC for Calbindin (a marker of PC) and BIP did not reveal obvious differences between *Dp2* mice and wild-type littermates (Supplementary Material, Fig. S1, and Fig. 4C, D). No significant apoptotic caspase 3 cells were observed in PCs of all mice (data not shown).

**Figure 4.**
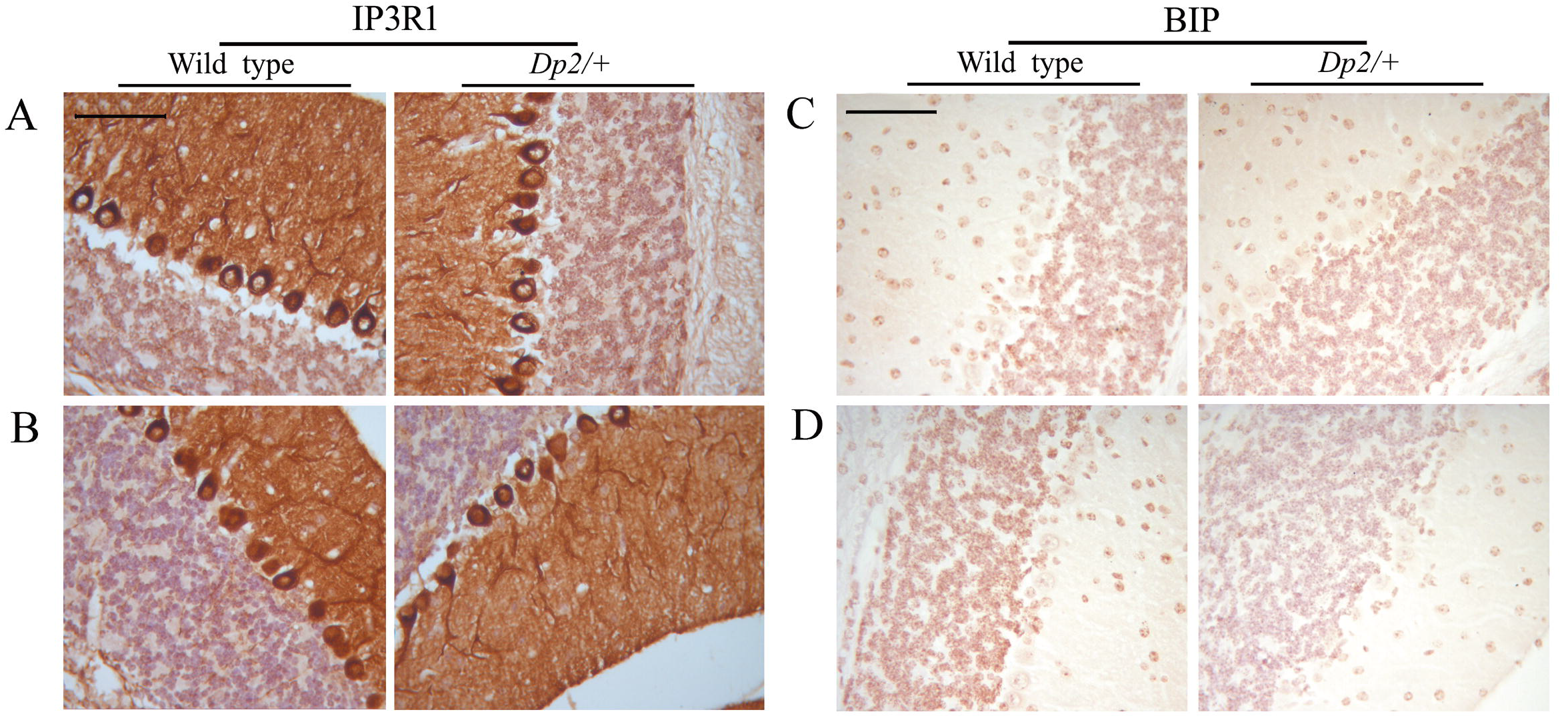
IHC analysis of cerebellum using anti-IP3R1 and anti-BIP antibodies. (A, B) IHC analysis of cerebellum using anti-IP3R1 antibody. *Dp2* mice that did not recover and littermate controls (A). *Dp2* mice that were recovered and littermate controls (B). The results showed the expression level of IP3R1 in the cerebellum of *Dp2* mice was equivalent to that of wild-type littermates. (C, D) IHC analysis of cerebellum using anti-BIP antibody. *Dp2* mice that did not recover and littermate controls (C). *Dp2* mice that were recovered and littermate controls (D). IHC analysis of cerebellum using anti-BIP antibody did not reveal obvious differences between *Dp2* mice and wild-type littermates. The expressions of IP3R1 and BIP are shown in brown. At least three mice were used for each analysis, and the representative data were shown. Scale bar=100 µm.

## DISCUSSION

The pupil is a hole located in the center of the iris of the eye. The iris is a contractile structure, consisting two layers: the front pigmented fibrovascular layer known as a stroma and, beneath the stroma, pigmented epithelial cells. The stroma connects to an iris constrictor muscle (pupillary sphincter) and a set of iris dilator muscle (pupillary dilator). Pupil size is controlled by the iris constrictor muscle and the iris dilator muscle, which act in opposition to cause miosis (constriction) or mydriasis (dilation) of the pupil in response to different levels of light. A greater intensity of light causes the pupil to constrict, thereby allowing less light in, to prevent damage to the retina from excess light (Davis-Silberman and Ashery-Padan, 2008). Hematoxylin and eosin staining showed an intact but short and unstretched iris, and the pilocarpine solution failed to constrict the mutant pupils in *Dp2* mice that did not recover. These results suggest that the *Dp2* phenotype is characterized by mydriasis, which is caused by a functional defect of the iris constrictor muscles.

Calcium ions influxes into the cytoplasm and their release from the endoplasmic reticulum (ER), allows actin-myosin interactions and contraction. One of the regulators of the intracellular Ca^2+^ is the IP3R1, which upon stimulation by the second messenger IP3 releases Ca^2+^ from the ER (Costa et al., 2010; Jiang and Stephens, 1994; Nakazawa et al., 2011; Sugawara et al., 2013; Walsh, 1994; Zhu et al., 2011). It is therefore reasonable to assume that the C1976Y mutation in IP3R1 affects Ca^2+^ release from the ER, which lead to contraction defect of the iris constrictor muscles.

Mice with *Ip3r1* heterozygous null mutations showed no mydriasis. These data suggest that the role of IP3R1 in iris constrictor muscle is tolerant of 50% residual channel activity. Given that IP3R1 forms a homotetramer, then only 1/16 assembled tetramers will contain four wild-type subunits. If a single variant subunit can block channel function, then 94% of tetramers will be non-functional, thus the *Ip3r1 ^C1976Y^* mutation in *Dp2* is likely to be acting by a dominant-negative effect.

The unfolded protein response (UPR) is a cellular stress response to prevent misfolded protein accumulation is the endoplasmic reticulum (ER), which strictly controls protein quality. However, prolonged induction of the UPR because of continuous expression and misfolding of mutant protein can lead to severe ER stress and induces cell apoptosis. This occurs in a variety of conditions with a genetic basis, including Alzheimer disease and cystic fibrosis. An ER chaperone, GRP78 (also known as the immunoglobulin heavy chain binding protein, BIP), acts as the master regulator of unfolded protein response (UPR) signaling to improve biogenetic processes and has a cytoprotective function against ER stress. BIP can interact with IP3R1 monomers and tethers them to ensure the fidelity of subunit assembly without stochastic misassembly (Mimura et al., 2007; Schroder and Kaufman, 2005; Wang et al., 2010). GRP78 overexpression significantly enhances IP3R-mediated Ca^2+^ release (Higo et al., 2010). Genetic studies have shown that the loss of BIP function leads to defective neural development and involuntary movement. Our study suggests that the unfolded protein response is activated by the *Ip3r1^C1976Y^* mutation, which leads to the recovery of pupil size in *Dp2* mice.

Mammalian normal ocular surface development involves a transient closure and reopening of the eyelid. In mice, eyelid re-opening takes place two weeks after birth (Geh et al., 2011; Tao et al., 2005). So, the functional defect of the iris constrictor muscles may not be harmful to mice in the first two weeks after birth. However, the fidelity of the subunit assembly of IP3R1 is important to PCs from an earlier period. Thus we can hypothesize that an unfolded protein response may be happen at an earlier period in PCs in *Dp2* mice.

## MATERIALS AND METHODS

### Mice

The *Dp2* mutant heterozygote mouse was generated via ENU mutagenesis using B6 mice. All animal protocols were approved by the Institutional Animal Care and Use Committee (IACUC) of the Yangzhou University Animal Experiments Ethics Committee with permission number SYXK (Su) IACUC 2017–0044, and carried out in accordance with the approved guidelines.

### Preparation of DNA and RNA

Genomic DNA was isolated from mouse tail tips by proteinase K digestion, phenol chloroform extraction, and ethanol precipitation. Total RNA was extracted from the brains of *Dp2* and wild type mice using TRIzol reagent (Invitrogen).

### Linkage analysis

*Dp2* heterozygotes of the B6 background were mated to D2 mice to generate F1 mice. Subsequently, F1 mice with dilated pupils were backcrossed with B6 mice to generate N2 mice. DNA samples from N2 mutant mice were screened by polymerase chain reaction (PCR) for microsatellite markers, with PCR products were separated on 4% agarose gels by electrophoresis. After refining the mutation location to a critical region, these genes in the region were then sequenced.

### Mutational analysis

The exons sequences (protein coding genes) or all sequences (non-coding RNA gene) of gene in critical region were amplified from *Dp2* heterozygotes and B6 RNA using RT-PCR and genomic DNA using PCR. Primer sequences are available on request. RT-PCR and PCR products were purified and sequenced.

### Histology

Eye and cerebellum samples were fixed at 4□ overnight. Fixed specimens were decalcified, dehydrated and embedded in paraffin wax, and 6-µm sagittal sections were obtained and H&E-stained using standard protocols.

### Immunohistochemistry

An antigen retrieval with citrate buffer was performed prior the antibodies incubation. The following antibodies were used: Anti-IP3R1 (1:400, ABCAM, Cat. ab5804), Anti-BIP (1:150, ABCAM, Cat. ab108615), Anti-Calbindin (1:400, ABCAM, Cat. Ab108404), Anti-Caspase 3 (1:100, CST, Cat.9579). Immunecomplexes were detected by using a biotinylated secondary antibody (1:400, Vector labs, Cat. BA1000) and ABC complex (Vector labs, Cat.PK6100) and AEC substrate kit (Vector labs, Cat. SK4100) or DAB substrate kit (Vector labs, Cat. SK4100), followed by brief counterstaining with Mayer’s hematoxylin.

## Acknowledgments

We would like to thank Paidashe Hove, Chathu Ranaweera and Ying Wang for critical reading of the manuscript. We are grateful for the assistance of Jianming Wang with photomicrography.

## Author contributions

Conceptualization: B.C., C.Q., L.C., Z.X.; Methodology: B.C., C.Q., L.C., M.D., Y.M., R.C., W.W., S.Y., H.W., X.D., M.G., Y.W.; Validation: B.C., C.Q., L.C., Z.X.; Formal analysis: B.C., C.Q., L.C.; Resources: B.C., Z.X.; Data curation: B.C., C.Q., L.C.; Writing-original draft: B.C., C.Q., L.C.; Writing - review & editing: B.C., C.Q., L.C., Y.W.; Supervision: B.C., C.Q., L.C., Z.X.; Project administration: B.C., Z.X.; Funding acquisition: B.C., Z.X.

## Funding

This work was supported by grants from the national natural Science Fund of China (31372269, 31000987), the Priority academic Program Development of Jiangsu higher education institutions (PAPD), and Yangzhou University Funding for Scientific Research (2016CXJ075).

**Supplementary information**

